# A Population Modelling Approach to Studying Age-Related Effects on Excitation-Contraction Coupling in Human Cardiomyocytes

**DOI:** 10.1101/2019.12.17.877514

**Authors:** Arsenii Dokuchaev, Svyatoslav Khamzin, Olga Solovyova

**Affiliations:** Institute of Immunology and Physiology UB RAS, Ekaterinburg, Russia; Institute of Immunology and Physiology UB RAS and Ural Federal University, Ekaterinburg, Russia

**Keywords:** Uncertainty quantification, Excitation-contraction coupling, Cardiac Electrophysiology, Sensitivity analysis, Computational modelling, Population of models

## Abstract

Ageing is the dominant risk factor for cardiovascular diseases. A great body of experimental data has been gathered on cellular remodelling in the Ageing myocardium from animals. Very few experimental data are available on age-related changes in the human cardiomyocyte. We have used our combined electromechanical model of the human cardiomyocyte and the population modelling approach to investigate the variability in the response of cardiomyocytes to age-related changes in the model parameters. To generate the model population, we varied nine model parameters and excluded model samples with biomarkers falling outside of the physiological ranges. We evaluated the response to age-related changes in four electrophysiological model parameters reported in the literature: reduction in the density of the K^+^ transient outward current, maximal velocity of SERCA, and an increase in the density of NaCa exchange current and CaL-type current. The sensitivity of the action potential biomarkers to individual parameter variations was assessed. Each parameter modulation caused an increase in APD, while the sensitivity of the model to changes in G_CaL_ and V_max_up_ was much higher than to those in the effects of G_to_ and K_NaCa_. Then 60 age-related sets of the four parameters were randomly generated and each set was applied to every model in the control population. We calculated the frequency of model samples with repolarisation anomalies (RA) and the shortening of the electro-mechanical window in the ageing model populations as an arrhythmogenic ageing score. The linear dependence of the score on the deviation of the parameters showed a high determination coefficient with the most significant impact due to the age-related change in the CaL current. The population-based approach allowed us to classify models with low and high risk of age-related RA and to predict risks based on the control biomarkers.

## 1 Introduction

Ageing is the dominant risk factor for cardiovascular diseases. A large body of experimental data has been gathered on cellular remodelling in the ageing myocardium from animals but very few experimental data are available on the age-related changes in the human cardiomyocyte. Mathematical models could be useful for predicting the adverse consequences of age-related changes in the myocardial function.

Recently, populations of human ventricular *in silico* action potential (AP) models were used to predict the clinical drug-induced arrhythmic risk of Torsade de Pointes (TdP) arrhythmia. The occurrence of cellular repolarisation abnormalities (RA) and the shortening of the electromechanical window (EMw), reflecting the difference between the durations of the electrical and mechanical systole, was evaluated in model populations and showed a high accuracy (close to 90%) in predicting TdP risk for a number of reference compounds [1, 2]. These *in silico* trials demonstrated a challenging promise of such analysis as a valuable tool for cardiotoxicity prediction.

Obviously, a similar approach can be used not only for drug safety testing but, more extensively, for assessing the pro-arrhythmic consequences of various interventions affecting the cellular mechanisms of excitation-contraction coupling.

The studies referred to above employed a surrogate index for drug effects on the electro-mechanical window evaluated by the effects of drug-induced modulation of the ionic currents on AP and Ca^2+^ transient duration because the cellular ionic models used in these studies do not allow one to evaluate possible direct drug effects on the contractile machinery. Such interventions could be simulated by integrating ionic models with cardiac contraction models and performing electromechanical simulations.

In this paper, we apply the experimentally calibrated population-of-models methodology [3, 4, 1, 2] o investigating variability in the response of cardiomyocytes to age-related changes in the parameters of the molecular and cellular mechanisms of excitation-contraction coupling. Besides, it is used for predicting the pro-arrhythmogenic effects of age-related cellular remodelling using human electro-mechanical cellular models developed by our research group [5].

The number of model parameters subjected to age-related modulations was restricted using several more or less standard approaches to model sensitivity analysis (SA) and uncertainty analysis (UA), such as Global Sensitivity Analysis and Polynomial Chaos Expansion [14]. This paper discusses in detail the use of these methods in rather wide parameter variation ranges and the need to account for possible implausible model outputs in the context of such variability.

Based on our model predictions, we used data analysis methods to suggest a surrogate model for estimating an individual risk of arrhythmia due to expected age-related parameter changes in a certain model from the control population. These results point to the potential of this approach for future possible implementation in personalized diagnosis of ageing-related cardiac abnormalities.

## 2 Methods

### 2.1 Electro-mechanical model of the human ventricular cardiomyocyte

In this study, we used a new combined electro-mechanical model (TNNP+M) of the human cardiomyocyte [5] based on the TNNP06 ionic model [7] of action potential (AP) and the “Ekaterinburg” model of mechanical activity and calcium handling in ventricular cardiomyocytes [8].

The model contains a biophysically detailed description of ion channels, pumps, and exchange currents, and also includes a detailed description of the concentration of intracellular sodium, calcium, and potassium. The electrical part of model mostly remains unchanged, the block describing the dynamics of calcium was modified by adding the equation for calcium troponin complexes (see Supplementary Material) linking the mechanical part with the electrical part of the model. The mechanical part of the model contains equations for myocardial stress, changes in the length of sarcomeres and the whole cardiomyocyte, including the representation of force-generating cross-bridges (Xb). The reader is referred to the Supplement for a detailed description of the model.

### 2.2 Control population of models

Using the methodology developed by B. Rodriguez group [3, 4, 1, 2], we constructed a control population of human ventricular electro-mechanical models based on the above TNNP+M model with a reference set of parameters. We varied randomly nine model parameters, having chosen, as suggested in [1], eight parameters for the conductances of main transmembrane ionic currents: fast Na^+^ current (G_Na_), transient outward K^+^ current (G_to_), rapid and slow delayed rectifier K^+^ currents (G_Kr_ and G_Ks_), inward rectifier K^+^ current (G_K1_), the L-type Ca^2+^ current (G_CaL_); and the Na^+^ – K^+^ (NKX) and Na^+^ – Ca^2+^ (NCX) exchangers (P_NaK_, K_NaCa_). The ninth parameter to be varied represented the maximal velocity of the SERCA pump (V_max_up_) as it affects the Ca^2+^ transient and subsequent mechanical activity in cells [9].

We randomly generated 20,000 parameter sets using the Latin hypercube sampling scheme with each parameter ranging from 0 to 200% of the reference value used in the baseline model. For each model sample, 200 cycles at a pacing rate of 1 Hz were computed to achieve a steady state. A set of several AP and Ca^2+^ transient biomarkers were calculated to compare model simulations with experimental data: AP duration at 40%, 50%, and 90% of repolarisation (APD_40_, APD_50_, and APD_90_); AP triangulation, defined as the difference between APD_90_ and APD_40_ (Tri_90_–_40_); maximum upstroke velocity (dV/dt_max_); peak voltage (V_peak_); resting membrane potential (V_rest_); and Ca^2+^ transient duration at 50% and 90% of repolarisation (CTD_50_ and CTD_90_). The population was then filtered by experimental biomarkers derived from the AP and Ca^2+^ transient recordings in isolated cells from undiseased human hearts as described in [2, 10].

In addition to this set of biomarkers, we also set min and max values of Ca^2+^ transient amplitude and the ratio between passive and active tension generated by myocardial preparations on the basis of data derived from experiments on animals of different species [11]. The complete list of biomarkers used in our study is given in Table 1.

**Table 1:** Experimental AP, Ca^2+^ transient and force twitch biomarker ranges used for calibrating the control population of human ventricular electro-mechanical *in silico* models, and extended biomarker ranges used for classifying non-implausible models in ageing populations.

| Control |  |  |  |  |  |
| --- | --- | --- | --- | --- | --- |
| Biomarker | Min Value | Max Value | Biomarker | Min Value | Max Value |
| $V_{\text{rest}}$ | -95mV | -80mV | $\text{CT}_{\text{peak}}$ | $0.5 \mu\text{M}$ | $2.5 \mu\text{M}$ |
| $V_{\text{peak}}$ | 10mV | 55mV | $\text{CTD}_{\text{peak}}$ | | 50 ms |
| $dV/dt_{\text{max}}$ | 100V/s | 1000V/s | $\text{CTD}_{50}$ | 65 ms | 420 ms |
| $\text{APD}_{40}$ | 85ms | 320ms | $\text{CTD}_{90}$ | 220 ms | 785 ms |
| $\text{APD}_{50}$ | 110ms | 350ms | $\text{FT}_{\text{d}}$ | | 10 |
| $\text{APD}_{90}$ | 180ms | 440ms | $\text{FT}_{\text{peak}}$ | | 150 |
| $\text{Tri}_{90-40}$ | 50ms | 150ms | $\text{FT}_{\text{peak}}/\text{FT}_{\text{d}}$ | 4 | 50 |
| Ageing |  |  |  |  |  |
| Biomarker | Min Value | Max Value | Biomarker | Min Value | Max Value |
| $\text{CT}_{\text{peak}}$ | | $3.0 \mu\text{M}$ | $\text{FT}_{\text{d}}$ | | 10 |
| $\text{CTD}_{90}$ | | 900ms | $\text{APD}_{90}$ | | 900ms |

Abbreviations: $V_{\text{rest}}$ , resting membrane potential; $dV/dt_{\text{max}}$ , maximum upstroke velocity; $V_{\text{peak}}$ , peak voltage; $\text{CT}_{\text{d}}$ , $\text{CT}_{\text{peak}}$ , $\text{Ca}^{2+}$ transient diastolic and peak values; $\text{FT}_{\text{peak}}$ , $\text{FT}_{\text{d}}$ , systolic and diastolic force during the twitch. $\text{APD}_{\text{XX}}$ , AP duration at XX% of repolarisation; $\text{Tri}_{90-40}$ , AP triangulation, defined as the difference between $\text{APD}_{90}$ and $\text{APD}_{40}$ ; $\text{CTD}_{\text{XX}}$ , $\text{Ca}^{2+}$ transient duration at XX% of decay. Both AP and $\text{Ca}^{2+}$ durations were computed starting from the instant of maximum $dV/dt_{\text{max}}$ .

The “control” population of virtual cardiomyocytes consisted of 240 model samples (Fig. 1) with biomarkers falling within the normal physiological range. The model acceptance rate in the final population was thus about 1.2% only. Such a low acceptance level reflects an essential role of the Ca^2+^ transient and mechanical biomarkers in the model filtration.

**Figure 1:**
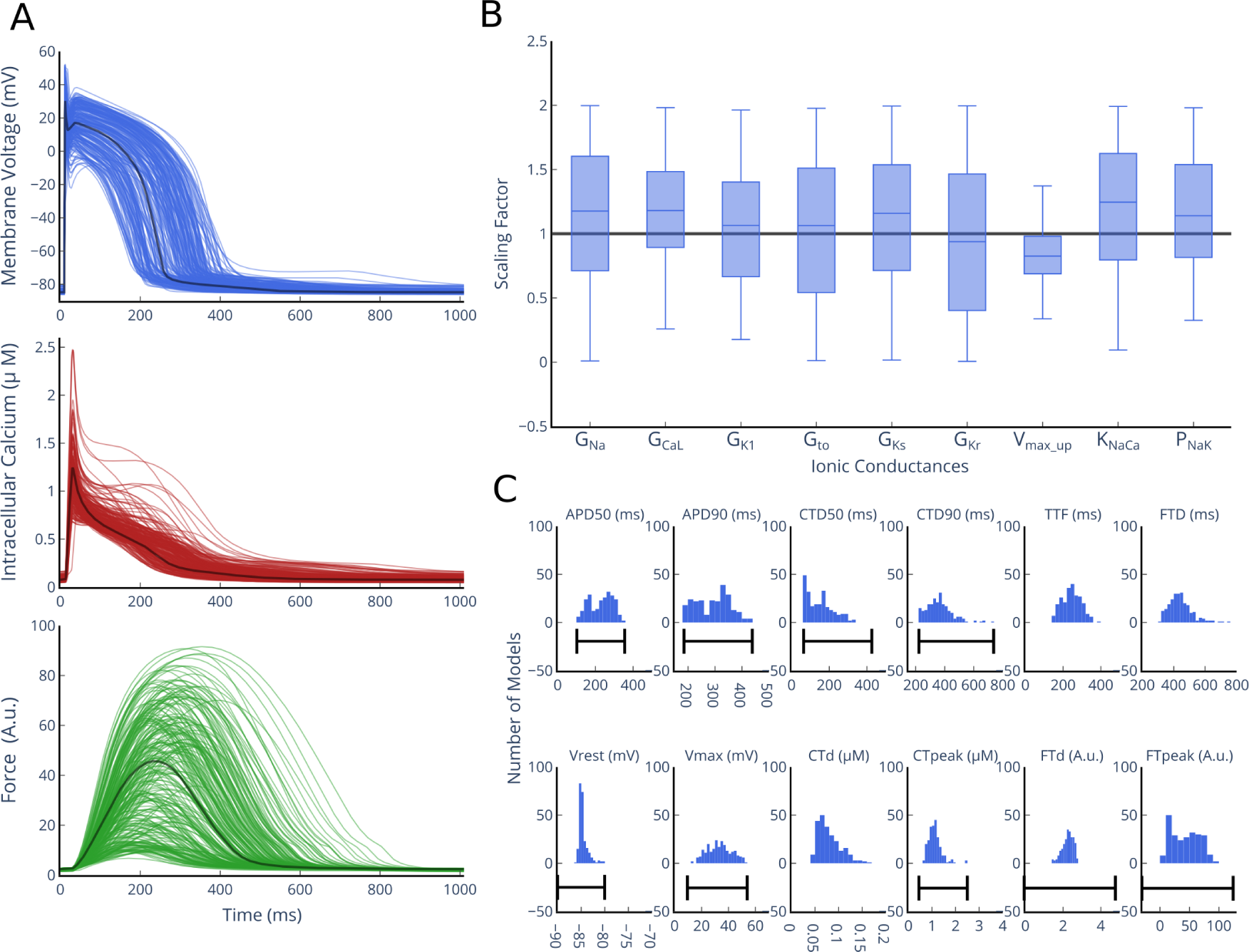
Population of human ventricular electro-mechanical models. (A) Traces of AP, Ca^2+^ ransient and Force twitch in the 240 models of the calibrated population. (B) Ionic parameter profile of the calibrated population, shown as scaling factors with respect to the baseline model. Central mark at box-plots corresponds to the median, box limits are the 25th and 75th percentiles, and whiskers extend to the most extreme data points. (C) Distributions of AP, Ca^2+^ transient and Force twitch biomarkers in the calibrated population (blue histograms) and experimental ranges (black lines): TPF, time to peak force; FTD, force twitch duration; other abbreviations as defined in Table 1.

The signals of of AP, Ca^2+^ transient and isometric force developed by every model in the population are shown together with the distributions of model parameters and biomarkers in Figure 1.

Abbreviations: V_rest_, resting membrane potential; dV/dt_max_, maximum upstroke velocity; V_peak_, peak voltage; CT_d_, CT_peak_, Ca^2+^ transient diastolic and peak values; FT_peak_, FT_d_, systolic and diastolic force during the twitch. APD_XX_, AP duration at XX% of repolarisation; Tri_90–40_, AP triangulation, defined as the difference between APD_90_ and APD_40_; CTD_XX_, Ca^2+^ transient duration at XX% of decay. Both AP and Ca^2+^ durations were computed starting from the instant of maximum dV/dt_max_.

All the simulations presented in this study were performed using Python3.6 language. Differential equations were solved using CVODE solver, a part of the open-source SUNDIALS suite [12] with just-in-time JIT Numba compilation to speed up calculation.

### 2.3 Model sensitivity analysis

To choose model parameters the modulation of which in ageing could essentially affect the electro-mechanical activity in the entire population, we performed an uncertainty quantification and sensitivity analysis of the baseline TNNP+M model accounting for uncertainty in the nine parameters varied in the population.

Variance-based global sensitivity analysis (GSA) was performed by computing the main and total Sobol sensitivity indices [13]. We used the Generalized Polynomial Chaos Expansion (gPCE) method [14], which allowed us to build a surrogate model and to compute Sobol indices analytically. The method is described in detail in the Supplement.

The mean decreasing accuracy (MDA) method was employed for sensitivity analysis as proposed in [16]. MDA had been originally introduced to identify feature importances in the Random Forest method [17]. Feature importance is estimated by measuring the quality of the prediction obtained from the classification/regression model after random permutation of features. In our study, the method was used as an alternative to Sobol sensitivity. Scikit-Learn package [18] was used for training/testing different machine-learning models and Chaospy Python 3.6 library for performing gPCE and calculating Sobol indices.

### 2.4 Aged population of models

The electro-mechanical response to age-related modulations in the four model parameters was evaluated against the control population. Three of the four parameters showed the most significant contribution to the model output biomarkers of AP, Ca^2+^ transient and cellular mechanics under consideration according to the model sensitivity analysis (see the Results for details). These were the density of NCX current (K_NaCa_), CaL current (G_CaL_), and the maximal velocity of SERCA (V_max_up_). The second consideration for choosing these three parameters was experimental evidence on the age-related remodelling in these ionic mechanisms in different animal species [26, 27, 28] suggesting their possible role in human ageing myocardium. Although the density of the potassium transient outward current G_to_ showed a moderate contribution to the model output biomarkers, we used it as well in this ageing population study because experimental data show its substantial change in ageing [25].

The direction of age-related parameter changes followed from the available experimental data suggesting a reduction in G_to_ and V_max_up_, and an increase in K_NaCa_ and G_CaL_ [21].

Two protocols of age-related change in the model parameters were performed. First, we gradually changed each parameter individually by 20, 50, 70% in every model of the control population to evaluate the sensitivity of the entire population to a parameter and to compare the contributions of the parameters to age-related abnormalities.

Then, we randomly generated 60 samples of the 4-variable vector of relative age modulations of the chosen parameters using normal distribution with 25% mean and 10% SD for every increment *δ*_k_, k = 1, …, 4, set as a percentage of the control parameter 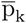. The age-related parameter values were computed as 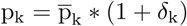. Root mean square deviation 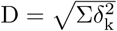 between the ageing and control parameters was used as a measure of model age-related remodelling in the parameter input space.

The distributions of each individual parameter deviation from the control and the distributions of the distance from the ageing parameter to the control are presented in Fig. S4 in Supplement. The mean distance between the ageing parameter vector and the respective control vector is close to 50% with a standard deviation of about 10% (54±9%), as expected from the above formula, indicting a two times greater distance for the vector with an average 25% modulation of each individual element set in the ageing population sample.

Each of the 60 age-modulated parameter sets was applied to every model from the control population, and, thus, 60 samples of the ageing populations and a total of around 15’000 age-related models were evaluated in the trial.

### 2.5 Arrhythmogenic score of age-related parameter modulation in the population of models

As in virtual arrhythmogenic drug testing [2], we checked the occurrence of the age-related effects on AP generation and Ca^2+^ handling associated with adverse events in the intact heart, particularly with the risk of TdP [19, 20].

We checked AP for repolarization and depolarization abnormalities (RA and DA, respectively), as in [1, 2]. Representative examples of age-related AP abnormalities are presented in Figure S4 in the Supplement.

For models not displaying AP abnormalities, a more than 6% increase in APD_90_: ΔAPD_90_ = (APD_90_aged_ – APD_90_)/APD_90_ > 6%, and a more than 10% decrease in the electro-mechanical window (EMw): ΔEMW = (EMw_aged_ – EMw)/EMw < −10% were considered as carrying a risk of adverse consequences (adverse events). Two EMw surrogates were computed. EMw_1_ = CTD_90_ – APD _90_ was used as suggested in [2]. Additionally, we computed EMw_2_ = FTD_p_ – APD _90_ as a more relevant measure of actual EMw, defined as the difference between the duration of cellular mechanical systole (time to peak force) FTD_p_ and APD_90_.

Moreover, we checked the model for implausible (ImP) outputs falling outside of the physiological ranges against the criteria provided in the bottom part of Table 1. Such ImP outputs were, as a rule, associated with an increase in the amplitude of the Ca^2+^ transient over an acceptable limit (see Table 1). Such outputs always caused abnormalities in force/shortening generation.

Every ageing model was paced at 1 Hz for 200 beats for each parameter modulation. The last trace of each simulation was compared with the corresponding control (without age-related modulation) and automatically checked for adverse events (RA, ΔAPD > 6%, ΔEMw < −10%, ImP).

A model was labelled as “Risky” if it had ΔAPD > 6% or ΔEMw < −10%, and models that did not fall into any of the adverse categories were labelled as “Norm”.

For each tested age-related parameter modulation (either individual parameter change or random samples of 4-parameter modulation) with distance D from the control, the arrhythmogenic score based on the frequency of adverse outputs (AO) was calculated as suggested in [1, 2] for virtual drug trials:

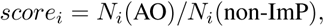

where *N*_*i*_ (RA) s the number of models with AO (either RA only or RA+ΔEMw_1, 2_) for the tested parameter modulation *i*, *N*_*i*_ (non-ImP) is the number of models in the population demonstrating non-implausible outputs.

We also assessed an index of “population loss” (PL) in the ageing population of models as a fraction of models with non-ImP outputs in the total population:

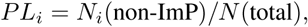

where *N*(total) is the total number of models in the control population subjected to ageing parameter modulation. The dependence of PL on parameter deviation from the control may be associated with a “survival curve” if an increase in parameter deviation is assumed to be related to age.

To assess the overall effect of ageing in 60 random ageing populations we used the following weighted scores accounting for adverse events in the same way as in [2, 1]:

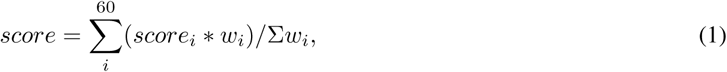

where *w*_*i*_ = 1/*D*_*i*_ is the weight inversely related to the tested distance *D*_*i*_. The overall score varies between 0 and 1, where 0 corresponds to a parameter modulation that does not provoke adverse events in any model from the entire population, while 1 corresponds to a parameter modulation that induces adverse events in 100% of the models.

## 3 RESULTS

### 3.1 A human *in silico* control population of electro-mechanical models

Figure 1 summarizes the behaviour of the control population of electro-mechanical models for human ventricular cardiomyocytes. The traces for AP, Ca^2+^ transient and isometric force twitch developed in the models are shown in the left panels (Figure 1.A). The boxplots for the distributions of the model parameters varied in the population are shown in Figure 1.B. The boxplots display a significant reduction in the parameter range as compared with the initial variation ranges due to the filtering out of implausible model samples. Moreover, both the medians of some parameters and the model traces are seen to significantly diverge from the reference values, e.g. most of the models demonstrate up-regulation of I_Na_ and I_CaL_ ionic currents (increased G_Na_ and G_CaL_), up-regulation of NCX and NKX exchangers, and down-regulation of I_Kr_ and V_max_up_. Figure 1.C displays histograms for the distributions of AP, Ca^2+^ transient (CT) and force twitch (FT) biomarkers versus the corresponding human experimental ranges (see also Table 1).

All the models demonstrate a normal AP phenotype, with all the AP and Ca^2+^ transient biomarkers falling within the experimental ranges.

### 3.2 Age-dependent input parameters in the model

For selecting model parameters to be subjected to age-related modulations, we performed an uncertainty and sensitivity analysis of the baseline model and identified parameters contributing substantially to the model output. As follows form the analysis of the first order Sobol sensitivity indices *S*_*i*_ for the nine varied parameters under consideration (for a heat-map of the indices see Fig. **??**, the first panel in the Supplement), three model parameters displayed the greatest contribution to the model output biomarkers of AP, Ca^2+^ transient and cellular mechanics. First, the density of CaL current G_CaL_ affects significantly the variance of AP temporal characteristics (up to 56% in APDs) and FT amplitude and duration (up to 54% of variation in FT_p_ and FTD). Second, the maximal velocity of SERCA V_max_ contributes more than 70% to CTD variance. Third, the density of NaK current P_NaK_ produces a major impact on the model outputs explaining 59% of variance in the diastolic Ca^2+^ level, and 41% and 39% variance in AP and [*Ca*^2+^]_*i*_ amplitude, respectively. Unexpectedly, most of the *K*^+^ conductances had no effect on the variation of the model outputs excepting G_K1_, which made a considerable contribution to the resting potential.

As noted in the Methods, the Sobol indices were computed for wide ranges of parameter variation. Therefore a great number of the model outputs fell outside of the acceptable ranges, contributing to potential overestimation of the model’s sensitivity as a result of accounting for an implausible parameter space.

Comparing the heat-maps for Sobol indices and MDA (see panels A and C in Fig. S3), we found that the significances of some parameters were different within the non-implausible space. For example, the model sensitivity of APDs to G_CaL_ decreased essentially below 10%, while sensitivity of APDs to P_NaK_ increased up to 70% in the non-implausible space. Moreover, variation of K_NaCa_ in the non-implausible space showed a notable contribution to the CTD biomarkers, though the Sobol coefficients for this parameter were close to zero.

At the same time, the list of parameters displaying the greatest contribution to APD, CTD, and FTD outputs remained the same, including G_CaL_, K_NaCa_, P_NaK_, and V_max_. Aging populations were simulated by three of these parameters: G_CaL_, K_NaCa_, V_max_ (see sub-Section C). The choice of these parameters was also supported by experimental evidence on their modulation in ageing (see the review [21]).

Although P_NaK_ demonstrated a high impact on the output biomarkers, we did not include this parameter into the ageing analysis because no direct experimental data were available on its change in ageing. In contrast, there are experimental data demonstrating the remodelling of I_to_ with ageing [26, 27, 28, 25]. The current I_to_ is known to contribute significantly to the AP shape, giving a spike-and-dome morphology. That is why we included *I*_*to*_ in the analysis of age-dependent consequences for AP phenotype.

### 3.3 Effects of individual modulation of age-dependent parameters

The effects of age-dependent increase in G_CaL_ and K_NaCa_, and decrease in G_to_ and V_max_ were tested against the control parameter values in the population. Figure 2 shows the evolution of the frequency of adverse events with increasing the age-related deviation of the model parameters from the control by varying individually each of the four parameters (top diagrams in panel A). Every ageing population was classified by observed adverse events into one of the following four classes: model outputs in the non-implausible range with no adverse events (Norm, green), ΔAPD_90_>6% and/or ΔEMw<−10% (Risky, blue tones), RA (orange), and implausible model outputs (ImP, rosy).

**Figure 2:**
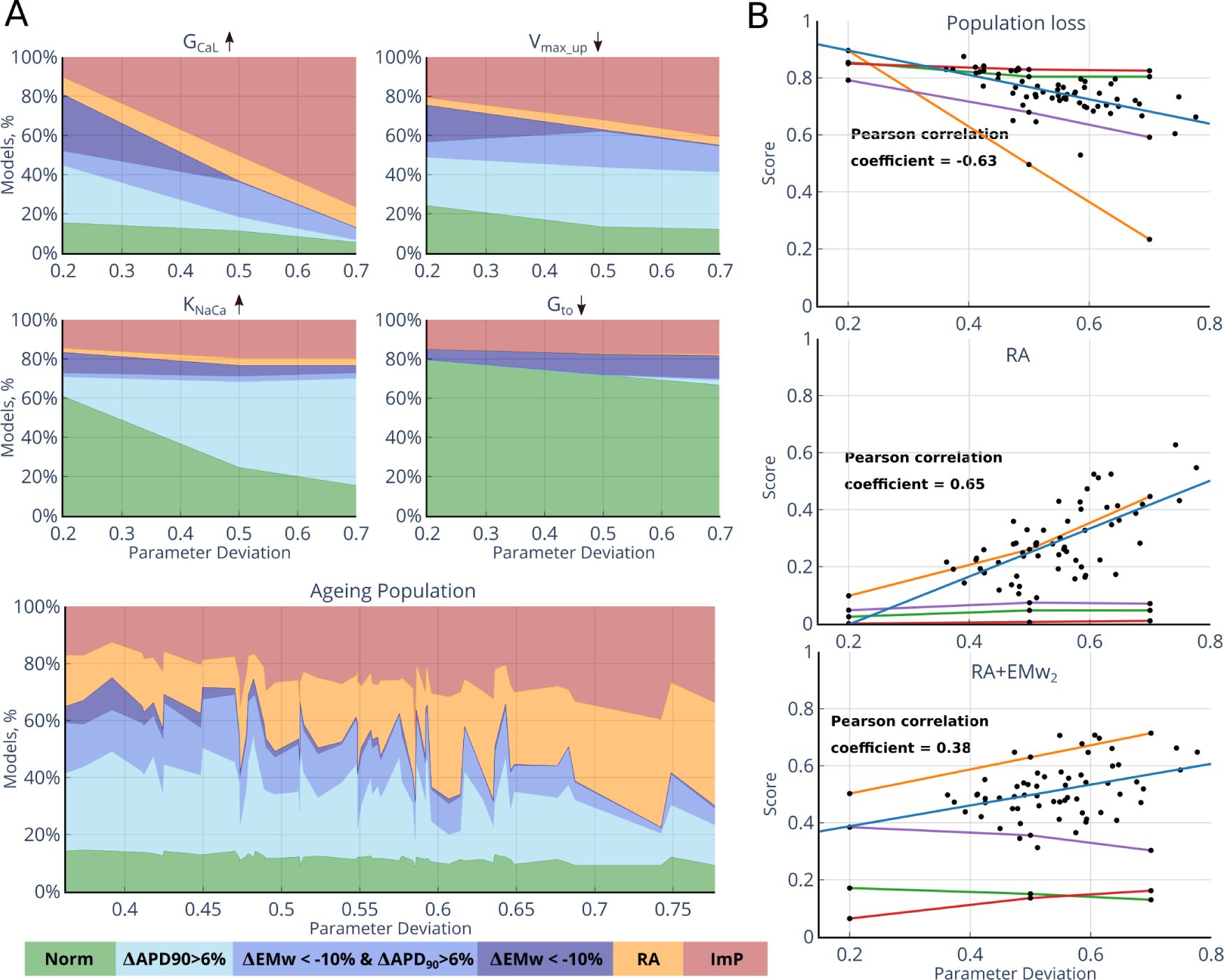
**A** Classification of adverse events during ageing. Colour maps indicate the “Norm” (green), “Risky” (blue tones), “RA” (orange), and “ImP” (rose) classes in the ageing populations. The “Risky” class contains models showing either ΔAPD_90_>6% or ΔEMw<−10% events (light and dark blue). Note that the ΔAPD_90_>6% and ΔEMw<−10% categories overlap if the events intersect (middle dark blue). EMw_2_=FTD_*p*_-APD_90_ was used as an indicator to demonstrate the frequency of ΔEMw<−10% events among the ageing populations (for similar diagrams computed for EMw_1_=CTD_90_-APD_90_ see Fig. S5). The top frequency diagrams show the results of age-related changes in individual parameters, the bottom diagram shows the classification of 60 ageing populations with four parameters varied. **B** Population loss and arrythmogenic score dependencies on age-related parameter deviation. The orange, green, red and purple lines show the dependencies obtained by individual modulation of G_CaL_, K_NaCa_, G_to_ and V_max_up_ respectively. The black dots are scatter plots for 60 ageing models with blue linear regression lines (p < 0.01).

Figure 2 clearly demonstrates that the overall frequency of adverse events increases with an increase in age-dependent parameter deviation from the control for each parameter tested. At the same time, the ratio of individual risk indicators in the overall frequency of adverse events is different for different model parameters. Although the frequency of ImP models in the ageing population increases with an increase in age-dependent deviation in every parameter, the increase in G_CaL_ has the highest impact on the occurrence of ImP in the model population. The fraction of ImP models and the slope of the “population loss” lines (Fig. 2.B, top panel) are significantly higher for G_CaL_ variation as compared to other parameter modulations. Whereas the fraction of RA events in the total population did not change essentially with an increase in G_CaL_, their frequency in the non-ImP parameter space increased with parameter ageing.

The fraction of “Risky” models with ΔAPD_90_>6% or ΔEMw<−10% events increased notably with increasing the deviation of K_NaCa_ or V_max_ from the control. An increase in K_NaCa_ causes the most notable increase in the fraction of ΔAPD_90_>6% models in the “Risky” category, while a decrease in V_max_ produces up to 20% of ΔEMw<−10% events. On the contrary, the fraction of “Risky” models in the total ageing population decreases with an increase in G_CaL_. As could be expected, decreasing G_to_ provokes fewer adverse events for any tested percentage as compared to the variation of the other parameters.

The middle and bottom panels in Figure 2.B show relationships between individual parameter deviations from the control and arrhythmogenic score defined as the frequency of adverse events based on the RA criteria only or combination of RA+ΔEMw<−10% as suggested in [2].

Increasing the age-dependent deviation of the parameters affects the scores differently. In line with the above analysis, the highest RA and RA+ΔEMw<−10% scores and the steepest age-dependence of the scores were obtained by increasing G_CaL_. For the RA risk indicator only, the contribution of the other parameters is considerably less pronounced, while the score based on the RA+ΔEMw_2_<−10% criterion is rather high in response to age-related decrease in V_max_up_.

This analysis allowed us to predict the greatest impact of these two parameters on the scores in the ageing populations for the deviation of the combination of four parameters (see Fig. 2, and sub-Section D).

To analyze the dynamics of model biomarkers that may be associated with enhanced vulnerability to arrhythmia in the ageing populations, we compared the distributions of APD_90_, CTD_90_, FTD*p* with those of the ΔAPD_90_ and ΔEMw biomarkers in the ageing models that did not produce any ImP or RA behaviour (Fig. 3).

**Figure 3:**
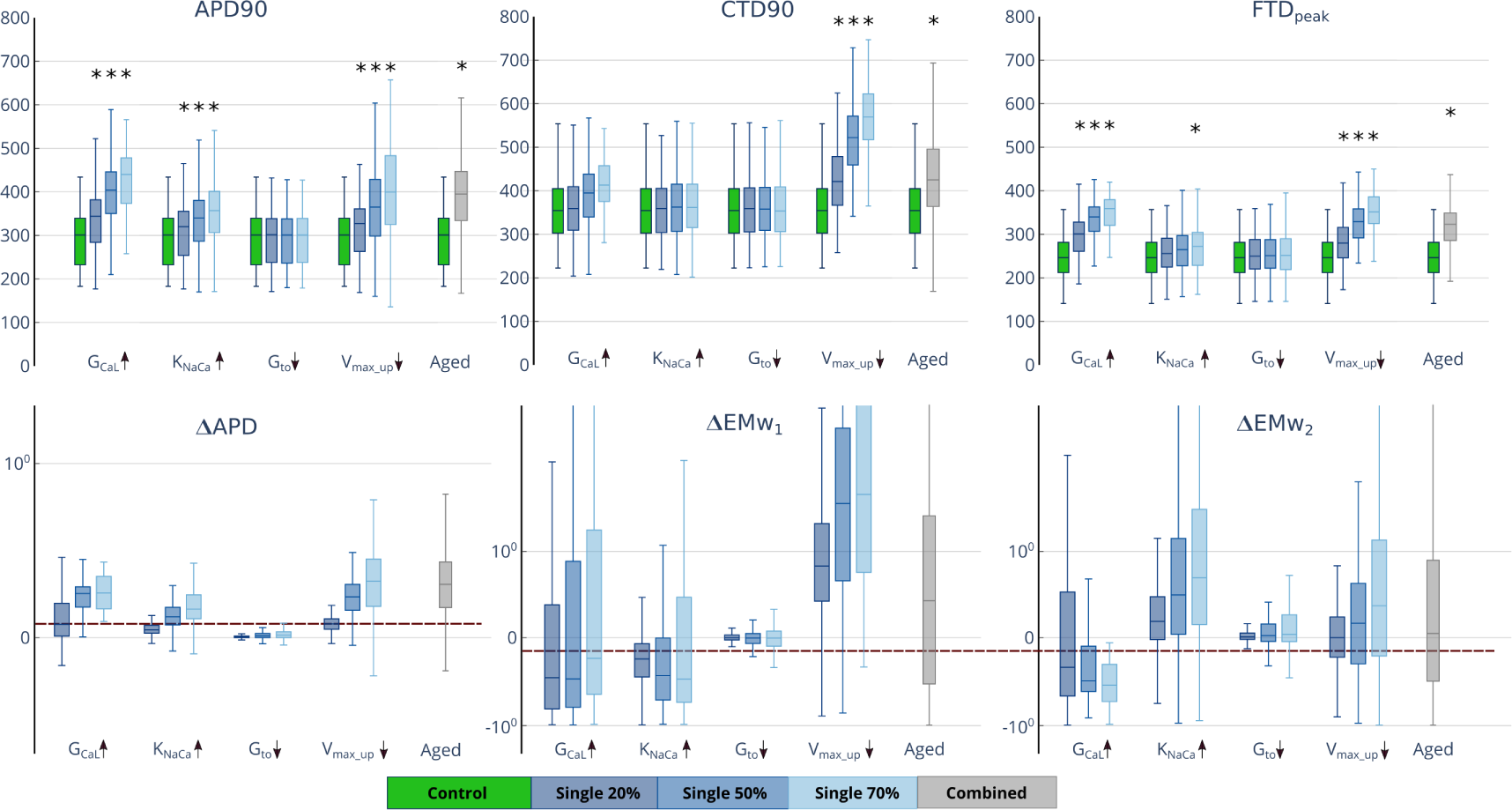
Age-related biomarkers. TOP panels show box-plots for the distributions of APD_90_, CTD_90_, FTD_*p*_ in model populations for age-dependent 20, 50, 70% deviations in individual parameters and in 60 aged populations with random parameter modulation. BOTTOM panels show box-plots for the difference between the biomarkers in the ageing and control models of the populations; ΔAPD_90_ and ΔEMw_1, 2_ are shown in the logarithmic scale.

Apparently, a decrease in G_to_ has no effect on the model output biomarkers. For other uni-directional parameter deviations (positive for G_CaL_ and K_NaCa_, and negative for V_max_), the effects on different biomarkers are different. First, an increase in the percentage deviation of each of the three parameters gradually increases the mean APD_90_ n the ageing populations with increasing the positive ΔAPD_90_. Mean ΔAPD_90_ vercomes the threshold level of 6% at 50 and 70% deviations of every parameter.

As could be expected, an age-dependent increase in parameter G_CaL_ causes an increase in the mean values of both CTD_90_ and FTD_p_ biomarkers (p<0.001). Although the average EMw_1, 2_ becomes shorter (mean ΔEMw_1, 2_<0 being above −10%), a large portion of ΔEMw_1, 2_ values in the ageing populations are positive and vary significantly in a wide range. This observation is consistent with a decrease in the frequency of ΔEMw_1, 2_<−10% events with increasing G_CaL_ as shown in Figure 2.A.

It is not surprising that even a steeper effect on CTD produces an age-related decrease in the parameter V_max_. This effect causes a significant increase in the mean CTD_90_ accompanied with an increase in the mean FTD_p_. At the same time, both mean ΔEMw_1, 2_ are positive and only a small portion of the models overcome ΔEMw_1, 2_<−10% indicator in the ageing population.

Although increasing K_NaCa_ has no effect on the mean CTD and FTD characteristics, it affects EMw_1, 2_ oppositely by shortening EMw_1_ with mean EMw_1_<−10%, but increasing EMw_2_ with mean EMw_2_>0.

Thus, although every parameter modulation predicts an arrhythmogenic increase in APD, inconsistent effects on ΔEMw_1, 2_ complicate the applicability of these indicators to the evaluation of the parameter contributions to the risk of adverse consequences of ageing in the model population.

### 3.4 Effects of combined modulation of age-dependent parameters

In the next series of simulations, we randomly varied all the four age-dependent parameters simultaneously as a random 4-dimensional vector (see Methods and Fig. S4 in Supplement). Sixty sampled parameter vectors were generated with an average parameter deviation (distance) D of 54±9% from the control (see Fig. S4) and corresponding 60 different ageing populations based on the control population of models were computed.

Figure 2 shows model classification for each of the 60 ageing populations, the scatter-plots of the arrhythmogenic score values and corresponding linear regression for the scores against the parameter deviation D. Figure 3 shows the overall distributions of APD_90_, CTD_90_, FTD_*p*_, and ΔAPD_90_, ΔEMw_2_ in all the ageing populations of models against the control values.

We found positive correlations between the age-related parameter deviations and the frequency of adverse events in the ageing populations with increasing arrhythmogenic scores in the non-implausible models of these populations (Fig. 2). The higher the age-distance, the higher the score suggesting an increase in the arrhythmogenic risk with cellular parameter modulation. The correlation coefficient for the score based on the RA criterion only is higher than that for RA+ΔEMw_2_ in the ageing populations (0.65 versus 0.38, p<0.01). This result reflects a non monotonous dependence of the score on individual parameter variations (Fig. 2) and high variability of ΔEMw_2_ with a positive mean value accompanied by an increase in the mean APD, CTD and FTD biomarkers (Fig. 3).

The regression lines for the scores in the random populations lie between the regression lines for populations with G_CaL_ and other parameters being varied, suggesting the highest contribution of age-dependent increase in G_CaL_ to the arrhythmogenic score.

To quantify the contribution of each parameter under combined parameter variation to the arrhythmogenic scores, we built linear models predicting the score value based on the deviations of the ageing parameters from the control (Table 2):

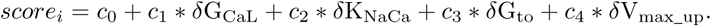

**Table 2:** Parameters of the linear model and determination coefficient (*R*^2^)

| Index | $c_0$ | $c_1$ | $c_2$ | $c_3$ | $c_4$ | $R^2$ |
| --- | --- | --- | --- | --- | --- | --- |
| RA | -0.21 | 1.19 | 0.18 | -0.02 | -0.51 | 0.89 |
| $RA + \Delta EMw_2$ | 0.27 | 0.87 | -0.12 | -0.02 | -0.13 | 0.86 |

The linear score models for both criteria (RA and RA+ΔEMw_2_<−10%) demonstrate similar accuracy (Table 2, see also Fig. S6 for correlations between computed and predicted scores, and Tables S1, S2 for ΔAPD and ΔEMw_1_ indicators in Supplement). As expected, the biggest coefficient in the linear model is associated with G_CaL_ deviation.

In the model for the RA score, the sign of the coefficients in the linear model is the same as that of parameter deviation meaning an increase in the score with increasing the absolute value of the deviation. The rather high coefficient for V_max_ deviation (although twice less than the coefficient of G_CaL_ shows a notable contribution of this parameter to the RA score. In the model for the RA+ΔEMw_2_<−10% score, the coefficient of K_NaCA_ deviation is opposite in sign to the deviation, meaning a decrease in the score with increasing K_NaCA_ deviation. This reflects a positive mean ΔEMw_2_ with its high variability under K_NaCa_ deviation.

To account for the higher severity of the arrhythmogenic risk for lesser parameter deviation D, we evaluated the overall arrhythmogenic scores of age-dependent parameter deviations using formula 1 (Table 3).

**Table 3:**
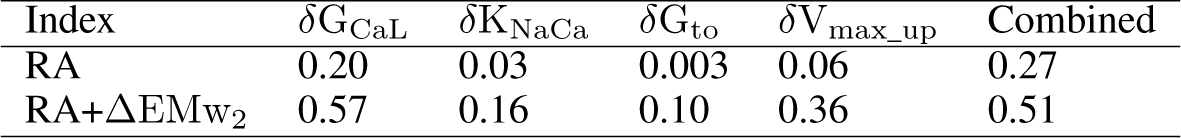
Overall scores under individual and combined age-related parameter modulation

| Index | $\delta G_{CaL}$ | $\delta K_{NaCa}$ | $\delta G_{to}$ | $\delta V_{max\_up}$ | Combined |
| --- | --- | --- | --- | --- | --- |
| RA | 0.20 | 0.03 | 0.003 | 0.06 | 0.27 |
| $RA + \Delta EMw_2$ | 0.57 | 0.16 | 0.10 | 0.36 | 0.51 |

It can be seen that the overall scores are higher for RA+ΔEMw_2_ than for the RA criterion alone, and the overall scores for the random ageing populations are close to those resulting from individual G_CaL_ modulations, suggesting its greatest contribution to the arrhythmogeneity in the ageing population of models.

### 3.5 Prediction of the RA score from control output biomarkers

We evaluated whether the AP, CT and FT output biomarkers from the control population of models may enable one to predict individual RA risks in ageing. For each model from the control population (an individual “young” subject), we computed individual probability of RA in ageing (“true” probability value) as the frequency of RA across the total number of ageing parameter samples applied to a given model (60 ageing samples tested in this study). It should be noted that if a model produced outputs falling outside of the non-implausible ranges, it was excluded from the regression analysis. Thus, the regression was built on 195 cases only. This gave us a data set on the relationship between the input control biomarkers (model features) and the risk of RA in ageing. Then the population was divided into two groups, a training and a test one, in the ratio of 70/30. The training data set was used for building a regression model of the RA probability (predicted probability value) using the extremely randomized trees algorithm [22] available in Scikit-Learn package. The regression model was tested by 5-fold cross-validation. The accuracy of the regression model constituted 0.85 and 0.77 for the training and test datasets, respectively. The scatter plot and linear regression of the predicted RA probability versus computed “true” values are shown in Figure 4.A. The ERT method allowed us to classify feature importances for the regression model (Fig. 4.B). Surprisingly, in our ERT model the first two most significant features proved to be force twitch amplitude and duration biomarkers, FT_*p*_ and FTD_*p*_, providing for about 40 and 15% of prediction accuracy, while APD_90_ and CTD_90_ give less than 5% each.

**Figure 4:**
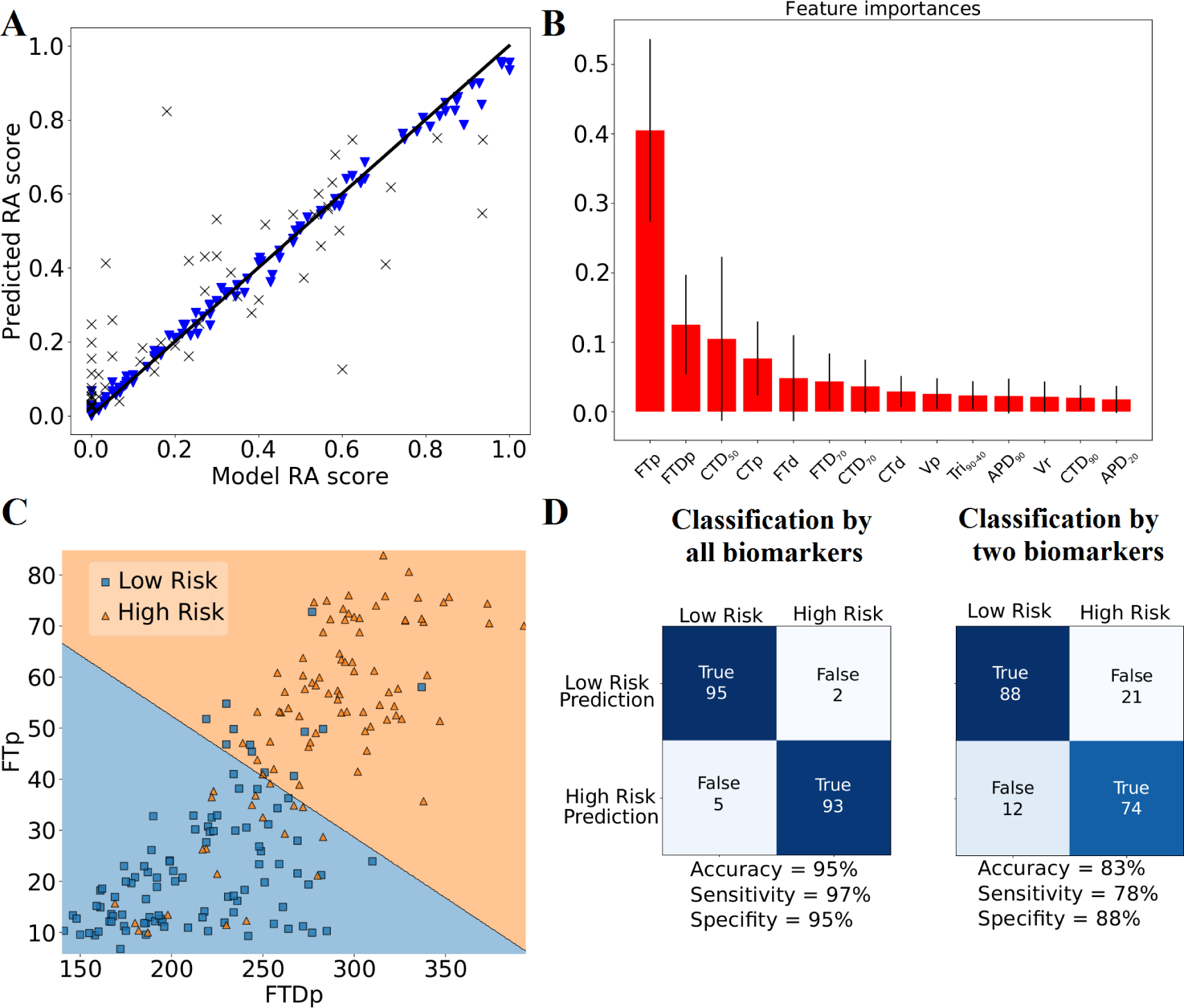
Prediction of RA risk in aging based on control model biomarkers. Panel A shows correlation between ERT predictions and computed RA probabilities in ageing. Blue triangles and black X labels show training and test model samples from the control population, respectively. Panel B shows the feature importance in ERT predictions. Panel C shows classification of the control population into low (<20%) and high (>20%) RA risk groups (blue and yellow) in the space of two most significant features, FTD_*p*_ and FT_*p*_. Panel D shows confusion matrices for the classification by all biomarkers and the two FTD_*p*_ and FT_*p*_ biomarkares. Columns show numbers of true and false predictions for the groups of low and high RA risk in the models from the control population with a corresponding classification accuracy, sensitivity and specificity to predict high RA risks.

Then we assessed the ability of the regression model to forecast low and high RA risks based on the ERT predictions of RA probability. First, we classified models from the control population according to the computed RA risk level. Models with true RA probability <20% were classified into the low-risk group, and those with higher true RA probability >20% into the high-risk group. The scatter plots of the two model groups in the control population are shown in Figure 4.C in the space of the two most valuable model features FT_*p*_ and FTD_*p*_. Comparing the ERT predictions of RA probability based on all the AP, CT and FT biomarkers with computed true values, we found the ERT model to have a 95 % accuracy accuracy in classifying low and high RA risks, with 97% sensitivity and 95% specificity in predicting high RA risks (Fig. 4.D, left). Finally, we assessed the ability of the classification model to forecast low and high RA risks on the basis of the two most valuable model features, FT_*p*_ and FTD_*p*_ only. The SVM classifier available in the Sklearn Package was used for separating the two groups in the 2-variable space as shown in Fig. 4.C. The linear separation model demonstrated an accuracy of 83% with a fairly rather high sensitivity at 78% and a high specificity at 88% in predicting high RA risks (see Fig. 4.D, right).

## 4 Discussion and conclusions

In this study, we developed a new simulation-based approach to predict the consequences of age-related cellular remodelling. A large number of ageing-related myocardial changes are already known, including ionic remodelling associated with changes in biophysical/biochemical mechanisms affecting the function of cardiomyocytes [21, 30, 31]. In this study, we focused on *in silico* assessment of electrophysiological consequences of age-related modulations in the cellular ionic parameters and on the prediction of the arrhythmogenic risk of cellular remodelling in ageing using human ventricular cellular models of electro-mechanical activity.

Cellular models have been extensively used as a valuable tool for studying different aspects of cardiac cellular physiology. They enable potential cross-links to be revealed in the regulation of the cellular function in norm and pathology and the role of cellular mechanisms in the response to interventions to be evaluated [32]. Along with the use of generic cellular models reproducing the average attributes of a population, a “population of models” approach has been actively developed recently [33]. It allows one to account not only for the natural variability in the cellular function even within one heart chamber but also for other sources of uncertainty inherent in mathematical models for biological systems.

This approach has been successfully applied to different research tasks such as model parameter identification [37], prediction of drug-induced cardiotoxicity [36, 35], and other challenges [33].

This study is based on the methodology developed by B. Rodriguez group and applied in human *in silico* drug trials for pro-arrhythmic cardiotoxicity prediction [1, 2]. This methodology allowed its authors to attain a high accuracy (close to 90%) in drug risk prediction for a great number of reference compounds they tested. The frequency of RA occurrence and EMw shortening were shown to be powerful indicators for predicting the clinical risk of arrhythmias based on ion channel information.

We made an attempt to use a similar methodology for predicting pro-arrhythmic effects of electrophysiological cellular remodelling in ageing. By way of extending this approach, our research group developed an electro-mechanical cellular model for evaluating the effects of age-related modulation of ionic mechanisms on the AP, CT and FT characteristics and proposed a more direct cellular surrogate of intact EMw as the difference between the mechanical systole (time to peak force FTD_*p*_) and excitation (APD_90_).

First, we developed a control population of electro-mechanical models. When we applied the same acceptable biomarker ranges as proposed in [2], a number of models that met the filtering conditions demonstrated implausible Ca^2^+ transient amplitudes and non-physiological mechanical features. This result suggests that in virtual trials of primarily electrophysiological issues under consideration (e.g. drug testing for pro-arrhythmic effects), special attention should be given to the features of simulated Ca^2+^ transient and contraction. We therefore imposed additional conditions on CT amplitude and the mechanical function of the cell based on the Frank-Starling law [11], and extended the acceptable limits for CTD_50_ based on the data from [10] (see Table 1).

Given these additional restrictions, the model acceptance rate dropped significantly from about 4000 models that were filtered by AP biomarkers only, to 1000 models that were filtered by AP and CT biomarkers, and finally to about 280 models that were filtered by AP, CT and FT biomarkers.

Supposedly, some molecular mechanisms that self-regulate AP, CT and FT outputs in the physiological ranges in healthy cells are not accounted for in the model, therefore the model formulations allow some outputs to overcome realistic limits. This means that some non-implausible model parameters could fall into a rather narrow space that is difficult to find using standard parameter identification methods applied to abstract mathematical systems. In this case, deletions of implausible cases from consideration is one of the suitable methods for limiting the non-implausible parameter space in the model [35].

In contrast to drug-testing studies, in which potential cellular targets are pre-determined and dose-response effects on the cellular ionic mechanisms are evaluated, we deal with a large uncertainty in the age-related effects on the parameters of electro-mechanical activity in human cardiac cells. While qualitative information is rarely reported, very few quantitative data are available for the age-related effects on cellular conductances and other mechanisms of excitation-contraction coupling. In these circumstances, we decided to choose only those parameters for subjecting to ageing modulations in the population of models whose contribution to the model outputs would be quantitatively substantial. To evaluate The contribution of the parameters to the cellular electrical and mechanical function was evaluated by the GSA method widely used in uncertainty analysis studies [13, 14, 15]. Note that this method is usually applied to parameter domains with rectangular borders. In our case, it would result in implausible parameters that would have to be accounted for in the Sobol indices. In order to restrict the sensitivity analysis to the non-implausible parameter space, we used the MDA method, which helped redistribute the roles of the model parameters in the variance of the model outputs. The same four most significant parameters, G_CaL_, V_max_up_, K_NaCa_ and K_NaK_, found using Sobol coefficients were also classified by MDA into the non-implausible parameter space. However, we did not include K_NaK_ into the age-related analysis because no direct data showing its possible modulation in ageing were found. Instead, we took G_to_ into consideration as the potential target of age-related parameter modulation based on the experimental data. However, further model analysis did not reveal any significant contribution of this parameter to age-related modulations of the model outputs in the ageing populations tested. Unexpectedly, the parameters of the K+ currents were not found to contribute significantly to the variance of the model outputs. This seems to be a specific feature of the TNNP ionic model, and a similar analysis has to be done using other ionic models combined with mechanical equations.

## 4.1 Arrhythmic scores and age-dependent changes

To simulate ageing populations, we randomly selected 60 sets of age-related parameter modifications by uni-directional deviation of the ionic parameters from the control values: G_CaL_ and G_NaCa_ were gradually increased in ageing, while V_max_up_ and G_to_ were reduced according to the available experimental data [26, 27, 28, 25]. These parameter modulations were applied to each of the models from the control population, and age-related changes in the model outputs were evaluated in the populations. It is still unknown what the molecular mechanisms of age-related conductivity changes are and how large the molecular changes can be. That is why we simulated parameter deviations in a rather wide range to be able to reveal the age-dependence of the effects.

Generally, our ageing models were able to reproduce ageing effects on cellular biomarkers observed in mammals, that is the average prolongation of AP, CT and FT [21, 31], reflected in an increase in mean APD, CTD and FTD in our ageing models (Fig. 3). This justifies our predictions of the molecular mechanisms we accounted when considering ageing-related remodelling in human cardiomyocytes.

It is well known that the risk of cardiac diseases increases significantly with age, including the risk of arrhythmias and sudden cardiac death [19, 20]. Several indices are widely used to assess the arrhithmogenic risk in the intact heart and in cardiac physiological models. One of those commonly used in drug testing is an increase in APD over a threshold level (in our study, the threshold is set at +6% according to [1]). The occurrence of RA events in isolated cells, in myocardial wedge preparations, or in the whole heart is common in the evaluation of pro-arrhythmic effects of different interventions. In addition, the shortening of EMw has been presented as an effective biomarker of pro-arrhythmia in several experimental studies [38, 39, 40, 41]. *In silico* drug trials using RA occurrence and frequency of RA+ EMw<−10% events provided a powerful methodology to predict the clinical risk of arrhythmias based on ion channel information [2].

Following these results by Passini and co-authors, we also used the frequency of adverse events as an indicator of arrhythmia risk in our ageing populations of models. In addition to the EMw surrogate based on CTD_90_ biomarker of Ca^2+^ transient as suggested in [1, 2] our second EMw estimate was based on the time to peak force FTDp computed from the force signal in the electro-mechanical model.

The score indices based on RA frequency alone and on the combined RA+ EMw<−10% criterion demonstrated positive correlations with the degree of age-dependent parameter deviations from the control (see Fig. **??**). However, the correlation was higher for the RA-only criterion, while the ΔEMw_1, 2_ indicators demonstrated very high variability in the ageing populations, different sensitivities to individual parameter deviations, and even opposite effects on ΔEMw_1_ and ΔEMw_2_ as in the case of K_*NaCa*_ age-dependent increase (Fig. 3). At the same time, the overall score accounting for the higher severity of adverse events at a shorter age-parameter distance from the control is twice greater for the RA+ΔEMw<−10% criteria that for RA only (Table 3), indicating its greater sensitivity to ageing parameter modulation. This result is consistent with the data of Passini et al [2], who found that the RA+ΔEMw<−10% score ensured a higher accuracy of drug-induced risk prediction at lower concentrations than the score based on RA alone.

It is important to highlight that drug effects on the ionic currents analyzed in [2] were induced by the inhibition of certain conductances, which reduced the corresponding ionic currents and subsequent increase in the APD_90_ and decrease in CT_90_ thus producing average EMw shortening. According to the authors, decrease in G_CaL_ as a major drug target was one of the essential reasons for CTD shortening, while drug-induced inhibition of K^+^ currents contributed substantially to APD prolongation. In contrast to drug effects on ionic current, in our study age-related effects on the ionic mechanisms are different for different currents, and one of the specific age-dependent parameter modulations is assumed to be increase in G_CaL_ conductance in accordance with experimental data [28]. At the same time, we did not modify the conductances of the essential K^+^ currents excepting G_to_, which had no much effect either on APD or on CTD.

The population of models approach allows one not only to predict the age-related risk to arrhythmia but also to evaluate the contribution of different ionic mechanisms to cellular vulnerability to age-dependent abnormalities. We compared the sensitivity of model populations to individual parameter changes (Fig. 2,3) and built linear models predicting risk scores from parameter deviations (Table 2). These results point to the most significant role of increase in G_CaL_ in age-dependent vulnerability to adverse events. Unexpectedly, the second most significant contribution to the pro-arrhythmic indicators came from decrease in V_max_up_ (see Fig. 2 and Table 2,3), which is apparently more important for the contraction rather than excitation phase.

### 4.2 Prediction of RA risk from the control biomarkers

The use of population of models approach in clinical applications requires the model to be able to predict pro-arrhythmic risk for an individual patient without knowledge of inherent cellular parameters, which are unlikely to be assessed in practice. To assess the RA risk for each individual model, we built a regression model based on biomarkers (features) from a control “young” population of models. Unexpectedly, the most significant features for predicting RA probability proved to be force biomarkers. We demonstrated that the use of just two biomarkers, force amplitude and time to peak, enabled us to predict RA risk with more than 80% accuracy in the ageing populations of models! This may be due to the fact that force biomarkers reflect not only free calcium in the cytosol but also calcium involved in the processes of cell contraction and intracellular homeostasis. Note that in this study we did not vary the mechanical parameters of the model, although it is known that the rate constants of contraction and relaxation decrease with age, myocardial stiffness increases, and the proportion of fast and slow myosine isoforms suffers changes [34, 31]. All these factors may affect significantly the characteristics of cellular contraction independently of the Ca^2+^ dynamics and, moreover, they may contribute to age-related mechano-dependent modulation of the Ca^2+^ transient and AP generation in cardiomyocytes.

These issues could be addressed using the methodology we developed in this study, which will make up the subject of our future research.

In conclusion, our main findings are as follows:

1. We developed a control population of electro-mechanical cellular models of human ventricular cardiomyocytes which can be used in the population based studies of the effects of different cellular interventions, including drug-testing, age-related remodelling, and cellular pathological remodelling.
2. The use of arrhythmogenic scores based on the RA and EMw criteria enable age-depending pro-arrhythmic risks of age-related parameter deviations to be predicted and classified.
3. Biomarkers of the myocardial contractile function constitute a powerful dataset for predicting the clinical risks of arrhythmias in ageing. Such information is often available from clinical data, and the integration of computer models in the existing pipelines for patient evaluation could improve diagnosis and personal treatment.

