## Supplementary Materials for "A Population Modelling Approach to Studying Age-Related Effects on Excitation-Contraction Coupling in Human Cardiomyocytes"

### Supplement

#### .1 Expanded Methods

##### .1.1 Model of excitation-contracion coupling

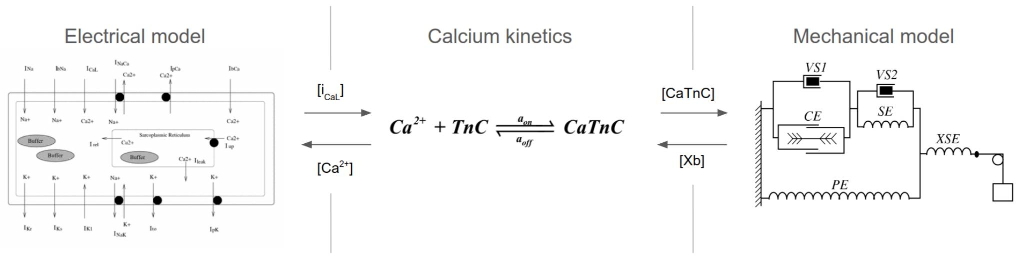


Figure S1: Full scheme of combined electro-mechanical model (TNNP+M) of the human cardiomyocyte [5] based on the TNNP06 ionic model [3] of action potential (AP) and "Ekaterinburg" model of the mechanical activity and calcium handling in ventricular cardiomyocytes [5].

The electro-mechanical TNNP + M model of a human ventricular cardiomyocyte [4] can be divided into mechanical and electrical parts. The electrical part describes the electrical function of a cardiomyocyte.The model contains a biophysically detailed description of ion channels, pumps, and exchange currents, and also includes a detailed description of the concentration of intracellular sodium, calcium, and potassium. The mechanical part of the model contains equations for myocardial stress, changes in the length of sarcomeres and the whole cardiomyocyte, including the representation of force-generating cross-bridges (Xb). It consists of a contractile element (CE), which is a generator of active force, three elastic (SE, PE, XSE) and two viscous (VS1, VS2) elements. The module of the mechanical activity has been developed by us earlier and used as a component in the electromechanical model 'Ekaterinburg – Oxford' [5] describing ECC in the cardiomyocytes of various animals (e.g. guinea pig and rabbit), adapted to each particular species via a parameter tuning. The mechanics of passive elastic and viscous elements, as well as the mechanical activity of the contractile element are described by a system of ordinary differential and algebraic equations we published earlier [5].

An important feature of this model is a mathematical description of the mechanisms of cooperativity: Xb-CaTnC, CaTnC-CaTnC and Tropomyosin end-to-end cooperativity [5]. Cooperative mechanisms of $Ca^{2+}$ activation of myofilaments (via kinetics of the CaTnC complex and force-generating crossbridges (Xb)) are the key feature of the model allowing it to reproduce a wide range of the effects of electro-mechanical coupling and mechano-electrical feedback in the myocardium.

The Ca-TnC kinetics is essential for initiation of the contraction, and therefore is described in the Ekaterinburg mechanical model by an ordinary differential equation separately from other intracellular $Ca^{2+}$ binding ligands. Moreover, the cooperative mechanisms of mechano-calcium feedbacks linking the Ca-TnC and Xb kinetics are described in the corresponding differential equations in the mechanical module of the Ekaterinburg model. These cross-links are the key mechanisms of cardiac ECC, which have to be reflected in our new model and will be used to evaluate a degree of mechano-calcium feedbacks and mechano-electrical feedbacks manifestations in the human cardiac cells.

Thus, in the combined TP+M model, we introduced a separate differential equation for Ca-TnC kinetics similar to that of the Ekaterinburg mechanical module and re-fitted the parameters in the algebraic quasi-stationary equation for the generalized calcium buffer of the TP model. In addition, we have changed some parameter values in the equation for the NCX current (iNaCa) from the TP model to fit the combined TP+M model to experimental data.

$$\frac{d[CaTnC]}{dt} = a_{on} \cdot ([TnC]_{tot} - [CaTnC]) \cdot [Ca^{2+}]_i - a_{off} \cdot e^{-k_A [CaTnC]} \cdot \Pi(Xb) \cdot [CaTnC] \tag{2}$$

where $TnC_{tot}$ is the total concentration of TnC in cytosol; $\Pi[Xb]$ is the cooperative dependence of dissociation of Ca-TnC complexes on the Xb concentration; $a_{on}$, $a_{off}$, $k_A$ are model parameters.

The following equation describes the time-dependent changes in free intracellular $Ca^{2+}$ concentration ($[Ca^{2+}]_i$):

$$\frac{d[Ca^{2+}]_i}{dt} = Ca_{i\_bufc} \cdot (\frac{I_{leak} - I_{up} \cdot V_{SR}}{V_c} + I_{xfer} - \frac{i_{bCa} + i_{pCa} - 2 \cdot i_{NaCa} \cdot C_m}{V_c \cdot F} - \frac{d[CaTnC]}{dt}) \tag{3}$$

Here, the first term describes Ca2+ release ($I_up$) and leakage ($I_leak$) from and $Ca_{2+}$ SERCA uptake ($I_up$) to the of sarcoplasmic reticulum ($V_c$ is the cytoplasmic volume, $V_SR$ is the SR volume). The second term ($I_{xfer}$) refers to the $Ca_{2+}$ diffusion leakage from the subspace into the cytoplasm. The third term describes $Ca_{2+}$ currents through the cell membrane (F is the Faraday constant, $C_m$ is the membrane capacitance). The descriptions of fluxes and ion currents are inherited from the TP model. The term $\frac{d[Ca^{2+}]_i}{dt}$ accounts for Ca2+ binding to TnC as provided by the Eq. 2.

Figure S2 A and B shows simulated force and length traces obtained under isometric conditions (i.e., fixed sarcomere lengths and transient increases in Ca, bold lines) and under isotonic loaded conditions at different afterloads (decreased from dark to light grey lines). Figure S2 C and F shows $Ca^{2+}$ and AP traces. The mechanisms of mechano-calcium and mechano-electric feedbacks reveal themselves in the prolongation of $Ca^{2+}$ transient and AP with decreased afterload. The $Ca^{2+}$ transient decay slows down via cooperative dependence of CaTnC kinetics on fraction of force-generated crossbridges. The slower $Ca^{2+}$ decay corresponds to the longer AP duration.

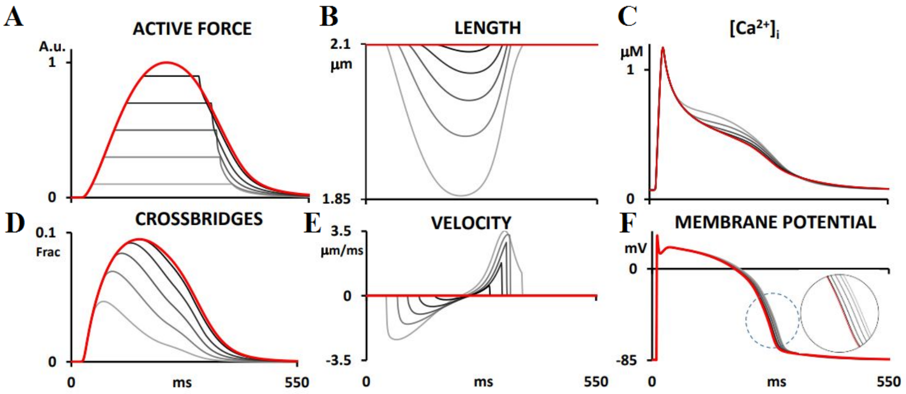


Figure S2: Afterload-dependent twitches. Bold (red) lines show the signals during isometric twitch. Grey lines show signals during isotonic contractions under various afterloads. The insert shows scaled-up load-dependent differences in the AP duration.

##### .1.2 Global Sensitivity Analysis

Sobol sensitivity indices provide fractional measures of the effect of uncertainty in each parameter on the resultant variance of the model output.

For model output, i.e. for a simulated biomarker $Y = f(\mathbf{X})$, where $\mathbf{X}$ is a vector of parameters (varying the conductivity of nine ionic channels and transporters). For the expectation and variance of $Y$: $\mathbb{E}(Y)$ and $\mathbb{V}(Y)$ the i-th first order Sobol sensitivity index is defined as:

$$S_{i_1} = \frac{\mathbb{V}(\mathbb{E}(Y|X_i))}{\mathbb{V}(Y)},$$

where $\mathbb{V}(\mathbb{E}(Y|X_i))$ is a conditional expectation, representing an expected value of the output $Y$ for a fixed value of input $X_i$. The i-th first order Sobol index can be interpreted as the fraction of the output variance that can be attributed to the variance of parameter $X_i$ alone (without interaction with other parameters). The common approach to performing GSA with the calculation of Sobol indices is Monte Carlo (MC) simulation due to its simplicity and easy implementation. However, it has an intrinsically low coverage rate which demands large amounts of model computations.

For UQ and GSA we used the Generalized Polynomial Chaos Expansion (gPCE) method [1], which allowed us to build a surrogate model and to compute Sobol's indices analytically.

The main idea of the gPCE methodology is that any stochastic output $Y$ can be spanned by multivariate orthogonal polynomials that are functions of the independent stochastic input parameters $\mathbf{X} = (X_1, ...X_k)$:

$$\mathbf{Y} \approx f_{gPCE}(\mathbf{X}) = \sum_{j=1}^{P-1} c_j \Psi_j(\mathbf{X}),$$

where $\Psi_j(\mathbf{X})$ are the orthogonal polynomial basis functions of parameter $\mathbf{X}$ of order $k$ and maximal degree $p$, $P = \binom{k+p}{p} = \frac{(k+p)!}{k!p!}$ is the number of unknown expansion coefficients $c_j$.

In our case $\Psi_j \equiv \Psi_{\boldsymbol{\alpha}^j} = \prod_{i=1}^{k} \psi_{\alpha_i^j}(X_i)$ in which $\psi_{\alpha_i^j}$ is a univariate Legendre polynomial of the order $\alpha_i^j$; $\boldsymbol{\alpha}^j$ is an integer sequence for the j-th multivariate polynomial: $\boldsymbol{\alpha} = \{\alpha_1, ..., \alpha_k\}$.

Due to orthogonality of the polynomial basis, the mean and variance of the response are $\mathbb{E}(Y) = c_0$ and $\mathbb{V}(Y) = \sum_{j=1}^{P-1} c_j^2 \mathbb{E}(\Psi_j^2(\mathbf{X}))$, and the gPCE derived Sobol indices are:

$$S_{i_1} = \frac{1}{\mathbb{V}(\mathbf{Y})} \sum_{\boldsymbol{\alpha} \in A_{i_1}} c_{\boldsymbol{\alpha}}^2 \mathbb{E}(\Psi_{\boldsymbol{\alpha}}^2), \tag{4}$$

For more detail description of method we refer the reader to [1, 2].

In addition to the standard Sobol sensitivity indices, we suggested a kind of hazard index of non-physiological model behaviour under parameter variation, which we refer to as model implausibility index (ImPI). We assumed that GSA can overestimate the model's sensitivity to parameter variation due to the accounting for a physiologically implausible input space. In other words, model parameter variation might strongly affect the biomarkers placing them outside of the physiological range due to abnormalities in repolarization and/or other adverse events. However, parameter variation in the non-implausible parameter space would not affect the biomarkers as much as predicted by the Sobol indices of GSA. To test this hypothesis, we constructed an ImPI for each biomarker Y analysed in this study.

It was computed as a relative deviation of the model output Y from either of the limits of acceptable biomarker range (if Y falls outside of the range) in a series of tested models with parameters being varied:

$$ImPI = \frac{\Delta Y}{RL}, \tag{5}$$

where $\Delta Y$ is the biomarker deviation from either minimal or maximal range limit ($RL$). Note that if the biomarker falls into the acceptable physiological range, $ImPI = 0$.

We performed sensitivity analysis for ImPI. Heat-map for the Sobol first-order sensitivity indices for ImPI function is shown in Figure S3.B.

In this study, we also used the Mean Decrease Accuracy (MDA) method for sensitivity analysis as proposed in [7]. MDA was originally introduced to identify feature importance in Random Forest method [8]. The importance of features is estimated by measuring the quality of the prediction of the classification/regression model after random permutation of features. Heat map for MDA indices is shown in Figure S3.C.

**MAIN FINDING FROM THE MODEL SENSITIVITY ANALYSIS**

According to the analysis of the first order Sobol sensitivity indices $S_i$ for the nine varied parameters in consideration (Fig. S3 **A**), the three model parameters displayed the greatest contribution to the model output biomarkers of AP, $Ca^{2+}$ transient and cellular mechanics. First, the density of CaL current $G_{CaL}$ essentially affects the variance of AP temporal characteristics (up to 56% in APDs) and FT amplitude and duration (up to 54% of variation in $FT_p$ and FTD). Second, the maximal velocity of SERCA $V_{max}$ contributes to more than 70% of CTD variance. Third, the density of NaK current $P_{NaK}$ produces an essential impact on the model outputs explaining 59% of variance in the diastolic $Ca^{2+}$ level, and 41% and 39% variance in AP and $[Ca^{2+}]_i$ amplitude, respectively.

Note that the sum of the first order Sobol indices over the nine parameters varied (the sum of elements from a column of the heat-map) for every model output is not equal to 1. Inequality of first Sobol indices to 1 reflects a significance of interactions between the model parameters in variation of the model output. The higher the total of the Sobol indices, the more independent the effect of each individual parameter on the output characteristic. For instance, the high total sum found for AP amplitude consisted of an obviously high 43% contribution of the density of fast $Na" + "$ current $G_{Na}$ and the second highest but not so clearly apparent 38% impact of $P_{NaK}$. Total sums of about 0.9 were also found for CTD biomarkers, with the major contribution of solely $G_{CaL}$.

Comparison of the Sobol heat-maps for model outputs and for MImPI (Fig. S3, panels A and B) demonstrates their great similarity, revealing that the same model parameters demonstrate high contribution to both model outputs and MImPI variance. This means that standard Sobol indexes cannot be used without accounting for restrictions on a non-implausible parameter space which is no structured and no rectangle as assumed for the method.

In this case, we used a more appropriate MDA method for the model sensitivity analysis in the no-structured non-implausible parameter space (Fig. S3, panel C). Comparison of the heat-maps of Sobol indices and MDA feature importance indices (Figure S3, panels A and C) shows similar contribution of most of parameters we evaluated to model outputs, including high impact of $V_{max}$ on CTD characteristics.

However, contribution of some parameters assessed by the Sobol and MDA indices is different. Density of NKX current $P_{NaK}$ shows higher MDA indices for APD biomarkeres (up to 50% for $APD_{90}$ and more than 65% for $APD_{40}$ and $APD_{50}$ variation). The model sensitivity of APD biomarkers to $G_{CaL}$ decreased essentially below 10% level, while sensitivity of CTD biomarkers to $P_{NaCa}$ increased up to 30% in the non-implausible space as compared to Sobol indices.

Unexpectedly, as shown by Sobol, ImPI and MDA indices, most of the $K^+$ conductances have no effect on variation of model outputs, except $G_{K1}$ which contributes to resting potential essentially.

**A**

| | $V_{rest}$ | $V_{peak}$ | $APD_{20}$ | $APD_{40}$ | $APD_{50}$ | $APD_{90}$ | $Tri_{90\text{-}40}$ | $CT_d$ | $CT_{peak}$ | $CTD_{peak}$ | $CTD_{10}$ | $CTD_{50}$ | $CTD_{70}$ | $CTD_{90}$ | $FT_d$ | $FT_{peak}$ | $FTD_{peak}$ | FTD | $F_{speed}$ | $FTD_{70}$ |
|---|---|---|---|---|---|---|---|---|---|---|---|---|---|---|---|---|---|---|---|---|
| $P_{NaK}$ | 0.057 | 0.392 | 0.105 | 0.087 | 0.068 | 0.058 | 0.241 | 0.591 | 0.417 | 0.018 | 0.057 | 0.044 | 0.087 | 0.127 | 0.045 | 0.384 | 0.266 | 0.344 | 0.022 | 0.362 |
| $K_{NaCa}$ | 0.031 | 0.037 | 0.1 | 0.047 | 0.028 | 0.003 | 0.125 | 0.019 | 0.0 | 0.021 | 0.007 | 0.032 | 0.023 | 0.001 | 0.001 | 0.014 | 0.003 | 0.016 | 0.001 | 0.015 |
| $V_{max\_up}$ | 0.0 | 0.01 | 0.007 | 0.032 | 0.052 | 0.125 | 0.187 | 0.032 | 0.028 | 0.448 | 0.664 | 0.745 | 0.756 | 0.746 | 0.093 | 0.013 | 0.145 | 0.109 | 0.022 | 0.075 |
| $G_{Kr}$ | 0.012 | 0.0 | 0.018 | 0.038 | 0.047 | 0.06 | 0.044 | 0.0 | 0.001 | 0.001 | 0.0 | 0.002 | 0.0 | 0.01 | 0.001 | 0.006 | 0.006 | 0.008 | 0.001 | 0.009 |
| $G_{Ks}$ | 0.01 | 0.0 | 0.025 | 0.062 | 0.066 | 0.035 | 0.001 | 0.001 | 0.001 | 0.001 | 0.0 | 0.002 | 0.0 | 0.008 | 0.001 | 0.006 | 0.008 | 0.009 | 0.001 | 0.01 |
| $G_{to}$ | 0.001 | 0.058 | 0.004 | 0.0 | 0.0 | 0.0 | 0.0 | 0.0 | 0.001 | 0.0 | 0.0 | 0.0 | 0.0 | 0.0 | 0.001 | 0.0 | 0.0 | 0.0 | 0.022 | 0.0 |
| $G_{K1}$ | 0.63 | 0.0 | 0.012 | 0.011 | 0.014 | 0.077 | 0.145 | 0.0 | 0.0 | 0.001 | 0.0 | 0.001 | 0.0 | 0.008 | 0.001 | 0.001 | 0.001 | 0.003 | 0.001 | 0.004 |
| $G_{CaL}$ | 0.098 | 0.012 | 0.387 | 0.553 | 0.56 | 0.49 | 0.129 | 0.244 | 0.2 | 0.12 | 0.123 | 0.041 | 0.013 | 0.01 | 0.113 | 0.54 | 0.524 | 0.404 | 0.022 | 0.424 |
| $G_{Na}$ | 0.0 | 0.428 | 0.153 | 0.013 | 0.005 | 0.001 | 0.008 | 0.0 | 0.001 | 0.0 | 0.0 | 0.0 | 0.001 | 0.001 | 0.001 | 0.0 | 0.0 | 0.0 | 0.001 | 0.0 |
| SUM | 0.839 | 0.938 | 0.811 | 0.843 | 0.841 | 0.849 | 0.88 | 0.887 | 0.649 | 0.608 | 0.852 | 0.867 | 0.879 | 0.91 | 0.256 | 0.965 | 0.953 | 0.895 | 0.092 | 0.899 |

**B**

| | $V_{rest}$ | $V_{peak}$ | $APD_{40}$ | $APD_{50}$ | $APD_{90}$ | $Tri_{90\text{-}40}$ | $CT_{peak}$ | $CTD_{50}$ | $CTD_{90}$ |
|---|---|---|---|---|---|---|---|---|---|
| $P_{NaK}$ | 0.022 | 0.022 | 0.007 | 0.019 | 0.001 | 0.121 | 0.411 | 0.017 | 0.11 |
| $K_{NaCa}$ | 0.001 | 0.022 | 0.0 | 0.0 | 0.004 | 0.15 | 0.0 | 0.115 | 0.006 |
| $V_{max\_up}$ | 0.022 | 0.022 | 0.0 | 0.001 | 0.085 | 0.131 | 0.024 | 0.63 | 0.357 |
| $G_{Kr}$ | 0.022 | 0.001 | 0.001 | 0.003 | 0.013 | 0.095 | 0.0 | 0.017 | 0.0 |
| $G_{Ks}$ | 0.001 | 0.001 | 0.007 | 0.007 | 0.006 | 0.006 | 0.001 | 0.013 | 0.003 |
| $G_{to}$ | 0.001 | 0.001 | 0.0 | 0.0 | 0.0 | 0.004 | 0.001 | 0.0 | 0.003 |
| $G_{K1}$ | 0.022 | 0.001 | 0.001 | 0.002 | 0.033 | 0.188 | 0.0 | 0.009 | 0.001 |
| $G_{CaL}$ | 0.001 | 0.001 | 0.104 | 0.124 | 0.154 | 0.081 | 0.191 | 0.076 | 0.012 |
| $G_{Na}$ | 0.001 | 0.022 | 0.001 | 0.001 | 0.0 | 0.035 | 0.001 | 0.001 | 0.0 |

**C**

| | $V_{rest}$ | $V_{peak}$ | $APD_{20}$ | $APD_{40}$ | $APD_{50}$ | $APD_{90}$ | $Tri_{90\text{-}40}$ | $CT_d$ | $CT_{peak}$ | $CTD_{peak}$ | $CTD_{10}$ | $CTD_{50}$ | $CTD_{70}$ | $CTD_{90}$ | $FT_d$ | $FT_{peak}$ | $FTD_{peak}$ | FTD | $F_{speed}$ | $FTD_{70}$ |
|---|---|---|---|---|---|---|---|---|---|---|---|---|---|---|---|---|---|---|---|---|
| $P_{NaK}$ | 0.029 | 0.129 | 0.348 | 0.683 | 0.669 | 0.512 | 0.105 | 0.0 | 0.257 | 0.043 | 0.058 | 0.073 | 0.09 | 0.158 | 0.215 | 0.341 | 0.163 | 0.086 | 0.368 | 0.106 |
| $K_{NaCa}$ | 0.205 | 0.027 | 0.075 | 0.088 | 0.08 | 0.045 | 0.2 | 0.0 | 0.131 | 0.412 | 0.272 | 0.128 | 0.099 | 0.034 | 0.044 | 0.073 | 0.031 | 0.071 | 0.073 | 0.088 |
| $V_{max\_up}$ | 0.035 | 0.01 | 0.021 | 0.023 | 0.03 | 0.08 | 0.159 | 0.0 | 0.196 | 0.31 | 0.415 | 0.695 | 0.735 | 0.707 | 0.04 | 0.057 | 0.381 | 0.449 | 0.051 | 0.374 |
| $G_{Kr}$ | 0.013 | 0.005 | 0.023 | 0.04 | 0.056 | 0.117 | 0.157 | 0.0 | 0.05 | 0.039 | 0.055 | 0.018 | 0.009 | 0.013 | 0.055 | 0.034 | 0.019 | 0.035 | 0.036 | 0.039 |
| $G_{Ks}$ | 0.013 | 0.005 | 0.018 | 0.071 | 0.09 | 0.088 | 0.044 | 0.0 | 0.038 | 0.014 | 0.034 | 0.015 | 0.011 | 0.013 | 0.058 | 0.039 | 0.024 | 0.028 | 0.04 | 0.032 |
| $G_{to}$ | 0.009 | 0.05 | 0.025 | 0.011 | 0.012 | 0.021 | 0.033 | 0.0 | 0.021 | 0.041 | 0.042 | 0.014 | 0.007 | 0.007 | 0.035 | 0.023 | 0.015 | 0.02 | 0.031 | 0.02 |
| $G_{K1}$ | 0.675 | 0.006 | 0.017 | 0.014 | 0.015 | 0.056 | 0.204 | 0.0 | 0.028 | 0.064 | 0.038 | 0.022 | 0.008 | 0.009 | 0.036 | 0.023 | 0.016 | 0.022 | 0.029 | 0.019 |
| $G_{CaL}$ | 0.011 | 0.013 | 0.128 | 0.023 | 0.024 | 0.048 | 0.038 | 0.0 | 0.255 | 0.051 | 0.052 | 0.022 | 0.032 | 0.049 | 0.481 | 0.39 | 0.334 | 0.271 | 0.35 | 0.297 |
| $G_{Na}$ | 0.01 | 0.754 | 0.343 | 0.047 | 0.025 | 0.032 | 0.059 | 0.0 | 0.024 | 0.027 | 0.034 | 0.013 | 0.008 | 0.009 | 0.036 | 0.021 | 0.017 | 0.019 | 0.024 | 0.024 |

Figure S3: **A** Heat-map of first order Sobol sensitivity indices $S_i$ for output model biomarkers at 9 varied input model parameters. **B** Heat-map of first order Sobol sensitivity indices $S_i$ for model Implausibility index ImPI (Equation 5) **C** Heat-map of MDA derived indices for biomarkers of non-implausible models at 9 varied input model parameters

#### .2 Supplementary Figures and Tables

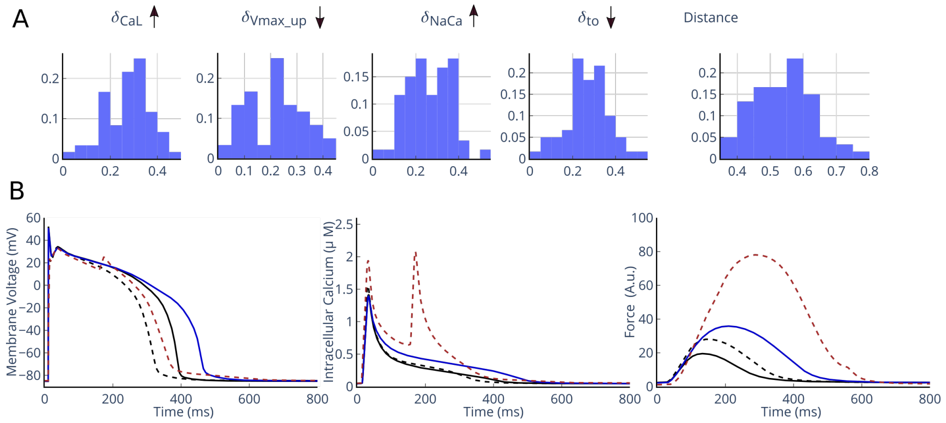


Figure S4: (**A**) Histograms of parameter deviations in the ageing model populations from the control and the distance D (see Methods Sec.) between them. Note that for $\mathrm{K_{NaCa}}$ and $\mathrm{G_{CaL}}$ the increments are positive (mean $\delta_k$ = 0.25) meaning an age-dependent decrease in the parameters; while for $\mathrm{G_{to}}$ and $\mathrm{V_{max\_up}}$ increments are negative (mean $\delta_k$ = -0.25) meaning age-dependent decrease in the parameters. (**B**) Examples of AP, $\mathrm{Ca}^{2+}$ and Force twitch transients in aging models demonstrating adverse events. Black lines show the control signals. Blue lines show a model with critically increased APD ($\Delta APD_{90} > 6\%$), red lines show an EAD model ($RA$) in the aging population.

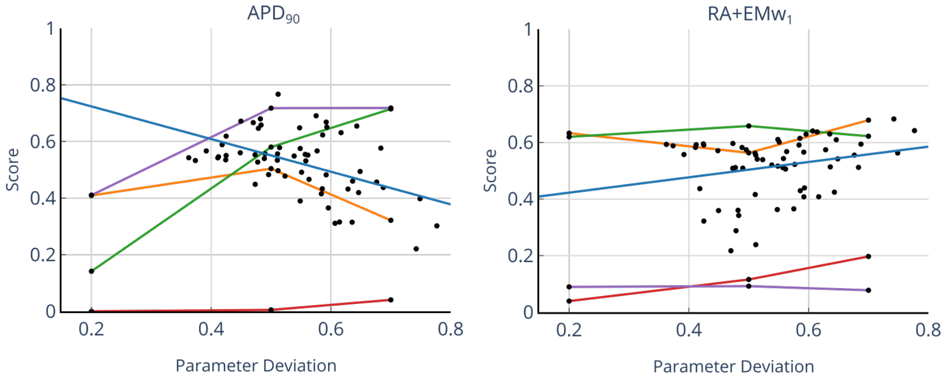


Figure S5: Arrythmogenic score dependencies on the age-related parameter deviation for $\mathrm{APD}_{90}$>6% and RA+ΔE$\mathrm{Mw}_1$<-10% criteria. Orange, green, red and purple lines show dependencies at $\mathrm{G}_{\mathrm{CaL}}$, $\mathrm{K}_{\mathrm{NaCa}}$, $\mathrm{G}_{\mathrm{to}}$ and $\mathrm{V}_{\mathrm{max\_up}}$ individual modulation, respectively. Black dots show scatter plots for 60 ageing models with blue linear regression lines. For $\mathrm{APD}_{90}$ based score the Pearson coefficient = -0.49, p<0.01, and slope k = 0.84; for RA+$\mathrm{EMw}_1$ based score correlation is not statistically significant (p =0.06 > 0.05)

Table S1: Parameters and determination coefficient ($R^2$) in linear models predicting the scores based on $\mathrm{APD}_{90}$>6% and RA+ΔE$\mathrm{Mw}_1$<-10% criteria: $score_i = c_0 + c_1 * \delta\mathrm{G}_{\mathrm{CaL}} + c_2 * \delta\mathrm{K}_{\mathrm{NaCa}} + c_3 * \delta\mathrm{G}_{\mathrm{to}} + c_4 * \delta\mathrm{V}_{\mathrm{max\_up}}$

| Index | $c_0$ | $c_1$ | $c_2$ | $c_3$ | $c_4$ | $R^2$ |
|---|---|---|---|---|---|---|
| $\mathrm{APD}_{90}$ | 0.90 | -1.06 | -0.06 | 0.02 | 0.28 | 0.81 |
| RA+$\Delta EMw_1$ | -0.21 | 1.19 | 0.18 | -0.02 | -0.51 | 0.89 |

Table S2: Overall scores at individual and combined age-related parameter modulation

| Index | $\delta\mathrm{G}_{\mathrm{CaL}}$ | $\delta\mathrm{K}_{\mathrm{NaCa}}$ | $\delta\mathrm{G}_{\mathrm{to}}$ | $\delta\mathrm{V}_{\mathrm{max\_up}}$ | Combined |
|---|---|---|---|---|---|
| $\mathrm{APD}_{90}$ | 0.42 | 0.34 | 0.008 | 0.54 | 0.53 |
| RA+$\Delta EMw_1$ | 0.62 | 0.63 | 0.08 | 0.09 | 0.51 |

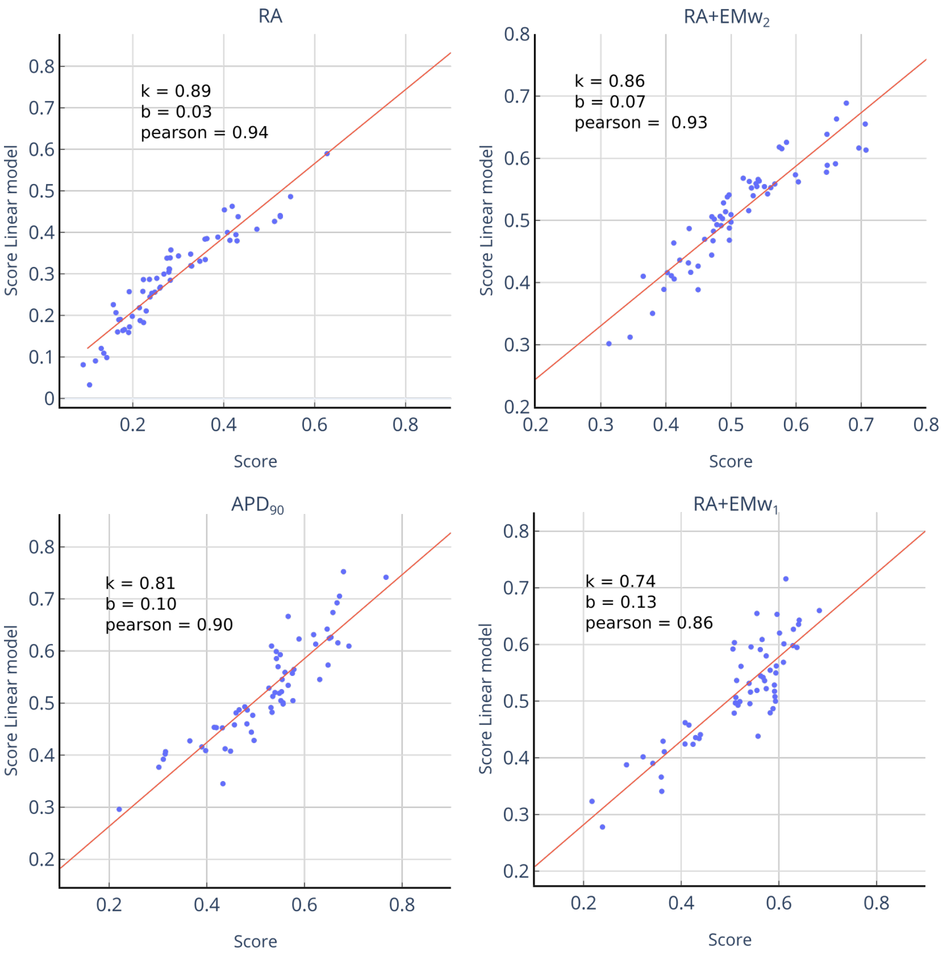


Figure S6: Correlation between the score values computed in the ageing populations and predicted from the linear model based on the deviations of aging parameters $G_{CaL}$, $K_{NaCa}$, $P_{NaK}$, and $\mathrm{V}_{\mathrm{max\_up}}$.

The linear model predictions show great correlation with Pearson coefficiten up to 0.94 and 0.93 for $RA$ and $RA + EMw_2$ based scores, respectively.

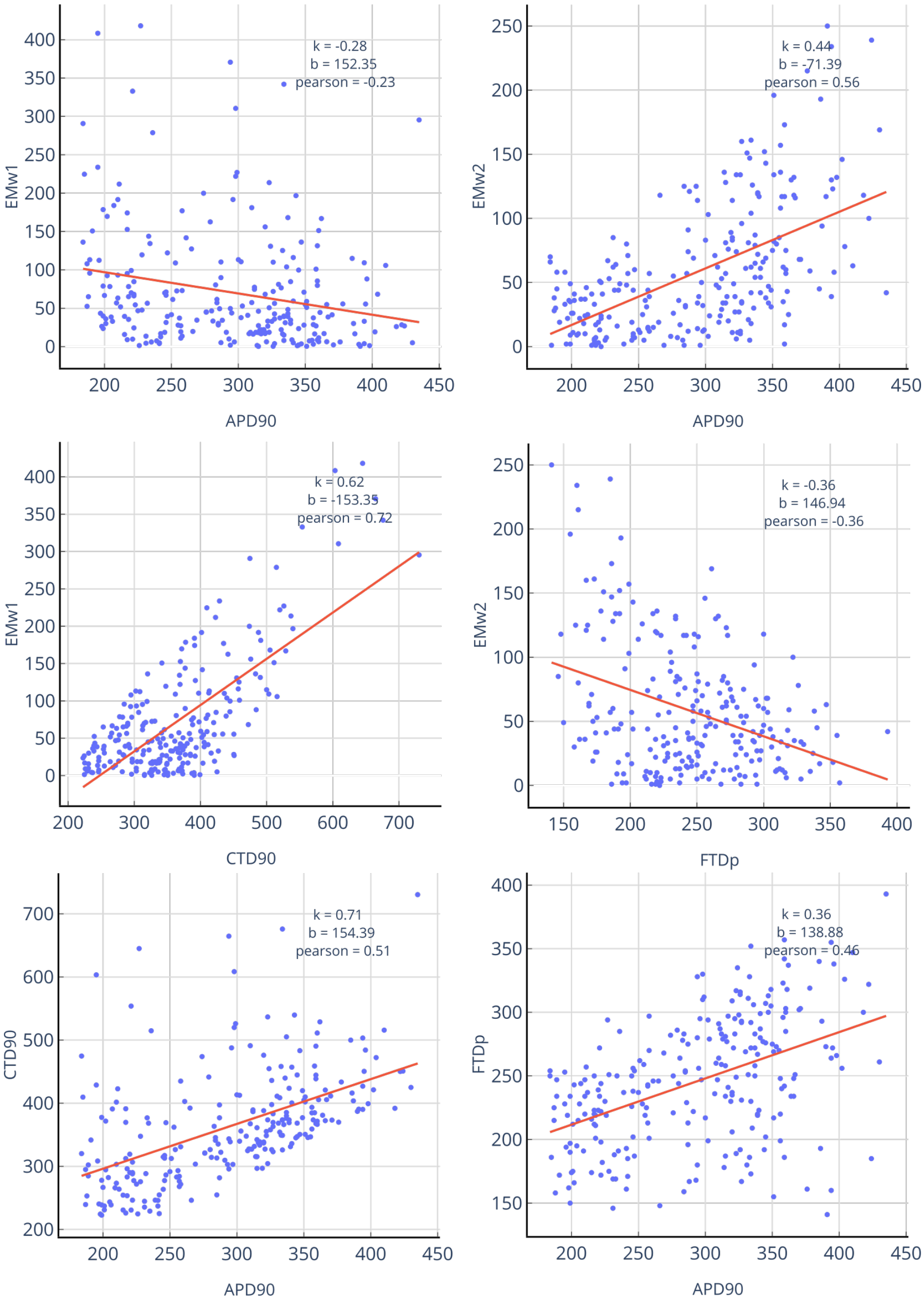


Figure S7: Relationship between APD, CTD, FTD, and EMw biomarkers in the 240 human ventricular models from the control population and their regression lines.

The scatter-plots show weak insignificant correlations between $EMw_{1,2}$ and $APD_{90}$, and between $EMw_{1,2}$ and $CTD_{90}$, $FTD_p$ respectively. In contrast to [6], weak correlations of EMw on either CTD or FTD characteristics, together with a high variability in the $\Delta EMw_1$ and $\Delta EMw_2$ in the ageing populations and different sensitivity to individual parameter deviations complicate applicability of these scores in the evaluation of ageing consequences for arrhythmogeneity.
